# Antibiotic treatment increases yellowness of carotenoid feather coloration in greenfinches (*Chloris chloris*)

**DOI:** 10.1101/2021.01.07.425726

**Authors:** Mari-Ann Lind, Tuul Sepp, Kristiina Štšeglova, Peeter Hõrak

## Abstract

Carotenoid plumage coloration is an important signal of quality, and plays an important role in mate choice in many bird species. However, it remains unclear what mechanism makes carotenoids an honest signal. Here, we test the hypothesis that carotenoid plumage coloration might indicate gut health. Parasitic and microbial infections can affect nutrient absorption due to decreased gut surface or by altered gut microbiome. We took an advantage of a naturally occurring coinfection of parasites inhabiting the upper and lower portions of the digestive track to distinguish between the direct and indirect effects of parasites on carotenoid acquisition. Protozoan coccidian intestinal parasites are widespread in greenfinches (*Chloris chloris*) and the majority of greenfinches are infected in nature. Trichomonosis is an emerging disease of the upper digestive track that causes high mortality among greenfinches. We captured wild greenfinches (N=71) and administered anticoccidial medication toltrazuril (TOLTRA) to one group, antibiotic metronidazole (METRO) that is also effective for treating *Trichomonas gallinea*, to the second group, and third group received no medication. In the METRO group, feathers grown during the experiment had significantly higher chroma of yellow parts, but there was no effect of TOLTRA on feather chroma. These results suggest that METRO increased the efficiency of carotenoid modification or deposition to the feathers rather than nutrient acquisition, and/or freed energy resources that could be invested in coloration. Alternatively, in accordance with shared pathway hypothesis, increase in efficiency of vital cellular processes might have occurred, as many microbial metabolites can modulate mitochondrial and immune function.

## Introduction

Carotenoid ornaments are important sexual signals that play a crucial role in mate choice in many vertebrate species, including birds. Males with brighter and more intense carotenoid feather coloration are preferred by females, are more likely to attain a mate, and invest more in taking care of nestlings (Hill, 1991). More colorful males live longer and have higher lifetime reproductive success (Cantarero et al., 2019). Honesty of sexually selected signals must be retained in order them to contain valuable information and avoid cheating (Zahavi, 1975), but after decades of studies, the mechanisms that guarantee honesty of carotenoid ornaments are still not clear.

Over the decades of research on carotenoid feather coloration, several non-mutually exclusive mechanisms have been suggested. For example, feather coloration has been shown to be associated with oxidative stress and antioxidant capacity (Alonso-Alvarez and Galván, 2011, Schantz et al., 1999, Tomášek et al., 2016). However multiple studies have failed to replicate these results (Sild et al., 2011, Mohr et al., 2019, Costantini and Møller, 2008) and metaanalyses have found only weak relationships between these variables and carotenoid coloration (Simons et al., 2012). It has also been proposed that carotenoids play an important role in mitochondrial membranes by participating in electron transport chain reactions and therefore signal organism’s potential to efficiently produce energy (the shared pathway hypothesis)(Hill, 2011). In other studies, carotenoids have been associated with resistance to parasites and parasite load (del Cerro et al., 2010, Hõrak et al., 2004). A recent meta-analysis found that metabolically converted carotenoids were related to parasite resistance and reproductive and parental quality, but unconverted carotenoids were not (Weaver et al., 2018).

Parasites can affect carotenoid coloration through compromised absorption of carotenoids in the gut. Many gut parasites cause inflammation of the gut surface that can reduce the absorption of carotenoids and overall energy acquired from the food (Tyczkowski et al., 1991). Among birds, protozoan coccidian parasites are widely distributed gut parasites that have been shown to affect digestive efficiency. Coccidia are intracellular parasites and their asexual and sexual multiplication inside of intestine’s epithelial cells damages animal’s ability to acquire nutrients from food (Joyner et al., 1975, Sharma and Fernando, 1975). In poultry, coccidian parasites cause extensive damage to the intestinal villi (Pout, 1967), inflammation of the gut surface (Sanches et al., 2020), leaky gut (Joyner et al., 1975), and reduced digestive efficiency (Sharma and Fernando, 1975, Russell Jr and Ruff, 1978). The damage caused by coccidiosis to intestines has been associated with malabsorption of carotenoids (Kouwenhoven and van der Horst, 1972, Ruff and Fuller, 1975, Tyczkowski et al., 1991) and other nutrients (Joyner et al., 1975, Sharma and Fernando, 1975). In captive greenfiches (*Chloris* chloris) goldfinches (*Carduelis carduelis*) and Nashville warblers (*Vermivora ruficapilla*) Isosporan coccidia caused thickening of intestinal walls (Swayne et al., 1991, Gosbell et al., 2020), which correlated negatively with weight (Gosbell et al., 2020). Coccidia have been shown to affect carotenoid coloration in wild birds (Hõrak et al., 2004, Baeta et al., 2008), but in the case of wild birds it is so far not clear if these effects occur through general condition decline or specific effect on gut integrity.

Other parasites, such as *Thrichomonas gallinae* and pathogenic bacteria, may not affect intestinal integrity directly, but decrease the amount of energy that is available to the individual from food. Protozoan *T. gallinae* is a parasite that causes lesions in the upper digestive tract in several species of birds, which in severe cases can block the esophagus and make the passage of food impossible (Amin et al., 2014). Avian trichomonosis caused by *T. gallinae* is historically common in pigeons and raptors, however since 2005 (Robinson et al., 2010) it has been reported as an emerging disease in greenfinches in Europe causing epidemics that have resulted in high mortality and decline of population of wild greenfinches (Robinson et al., 2010, Chavatte et al., 2019). Avian trichomonosis is an example of a relatively recent event where a parasite spills over to another taxonomic group with drastic effects to a new host species (Robinson et al., 2010).

Although several studies in poultry have detected the effect of coccidian parasites on decreased nutrient absorption and carotenoid levels, few studies have considered this mechanism in coloration of wild birds. In contrast to poultry, where most economically relevant coccidian parasites are from genus *Eimeria*, predominant coccidian genus in many wild species is *Isospora*, which might have different effects on the host. However, some studies in wild birds have revealed similar results to poultry. Experimental infection of greenfinches with coccidian parasites Hõrak et. al (Hõrak et al., 2004) showed parallel patterns of decreased carotenoid, serum albumin and triglycerides in serum, and decreased carotenoids in the feathers of greenfinches, which points to reduced digestive and absorption capacity. Infection with *Isospora sp.* coccidians increased fat content in the feces of greenfinches (Meitern et al., 2016). Carotenoids are fat soluble and carotenoid deficiency and malabsorption of lipids are likely related (Surai et al., 2001). In house finch (*Haemorhous mexicanus*) redness of carotenoid coloration correlated with digestive efficiency - individuals with redder ornaments absorbed more fats from their diet (Madonia et al., 2017). The effect of other infections, such as trichomonosis, on carotenoids acquired through diet has not received much attention. However, in greenfinches it has been shown that melanin, but not carotenoid coloration, predicted mortality, and black tail feathers became darker after the emergence of trichomonosis, but no change was observed in the yellow feathers (Hõrak and Männiste, 2016). Therefore, different parasites might induce selection pressure on different components of bird coloration.

The goal of this study was to test whether treatment with two different anti-parasitic medications affects plasma carotenoid levels and carotenoid-based feather coloration in greenfinches. Chroma of yellow parts of the feathers is related to carotenoid content in the feathers of greenfinches (Saks et al., 2003). Greenfinches in our study population are naturally infected by coccidian parasites (*Isospora spp.*) (Sepp et al., 2012). We have previously detected characteristic symptoms of trichomonosis in some birds that have died in the aviary (Männiste and Hõrak, 2014). We treated the birds with anticoccidial drug Toltrazuril (TOLTRA), which is designed specifically for treatment of coccidiosis and lacks known effects on microbes other than apicomplexans (Hackstein et al., 1995), and the other group received nitroimidazole group antibiotic, metronidazole (METRO), that targets protozoan parasites of trichomonas genus and wide spectrum of anaerobic bacteria (Amin et al., 2014).

As carotenoids have been associated with signaling overall parasite resistance (Weaver et al., 2018) we hypothesized that medicating birds with TOLTRA and METRO results in higher plasma carotenoids and higher chroma in lab-grown feathers in both groups compared to the control group. Our study design allows us to distinguish between the effects that parasites have on intestinal health and therefore nutrient absorption, vs general health state and/or metabolic processes that enhance deposition of carotenoids into the feathers. Accordingly, we predicted that if intestinal integrity is the key factor linking intestinal parasite infection to carotenoid coloration, treatment with either of the drugs (TOLTRA or METRO) affects plasma carotenoid concentration. Alternatively, if the key factor is overall health state, then birds treated with METRO show greater increase in plumage carotenoid coloration.

## Methods

### Study system

Wild male greenfinches (N=71) were captured in mist nets at bird feeders in a garden in the city of Tartu, Estonia (58°22′N; 26°43′E) on 5th, 6th, and 8th January 2015. Greenfinches are gregarious medium-sized (c. 28 g) seed-eating passerines native to the western Palearctic region (Cramp and Perrins, 1994). Males are more colorful (Svensson, 1992) with yellow, carotenoid-based (Stradi et al., 1995) markings on the sides of the tail feathers, primaries, primary covers and breast, while females lack full yellow tints in their plumage (Cramp and Perrins, 1994). Greenfinches incorporate two main carotenoids - canary xanthophylls A and B - into feathers to develop yellow color. Canary xanthophylls A and B are metabolically converted from dietary lutein and zeaxanthin (Stradi, 1998, McGraw et al., 2002). The birds were housed indoors in individual cages (27 × 51 × 55 cm) with sand-covered floors in a single room where they had visual contact with their neighbors. The average temperature in the aviary during the experiment was 15.5 ± 1.0° (SD) °C, and average humidity was 57 ± 7 (SD) %. The birds were supplied *ad libitum* with sunflower seeds and tap water and were exposed to a natural day-length cycle using artificial lighting by luminophore tubes. The birds were released back to their natural habitat on 3rd March 2015. The study was conducted under license from the Estonian Ministry of the Environment (Licence # 1-4.1/11/100, issued on 23rd March 2011), and the experiment was approved by the Committee of Animal Experiments at the Estonian Ministry of Agriculture (decision # 95, issued on 17th January 2012)

Birds were divided into three approximately equal-sized groups based on similar age composition (yearlings vs. older, determined based on plumage characteristics), body mass, and coccidian infection intensity, recorded on 15th January. Timeline of the study is depicted on Fig. 1. On the evening of 19^th^ January, the birds in two groups subjected to medication treatment started to receive either TOLTRA (24 birds) or METRO (24 birds) with their carotenoid-enriched drinking water. Twenty-three control birds received just carotenoid-enriched water. Birds in the anticoccidial medication group received 2 ml/L solution of Intracox Oral (Interchemie, Castenary, the Netherlands), containing 25 mg/L toltrazuril. METRO (Fresenius Kabi Polska, Kutno, Poland) was administered in concentration of 400 mg/L. Both drugs were dissolved in carotenoid solution (1 ml/L mix of lutein and zeaxanthin (20:1, w/w), prepared from OroGlo brand 15 Liquid Pigmenter with 15 g/kg xanthophyll activity (Kemin AgriFoods Europe, Herentals, Belgium)). Carotenoids were added to drinking water to compensate for naturally low carotenoid content of sunflower seeds. Medication lasted for 10 days, and carotenoid supplementation lasted until the birds were released. Birds were weighed and blood sampled on 19^th^ January and 30^th^ January in order to record the effects of treatments on plasma carotenoid concentrations. Blood sampling of birds took place in the mornings before the lights turned on.

**Fig. 1.**
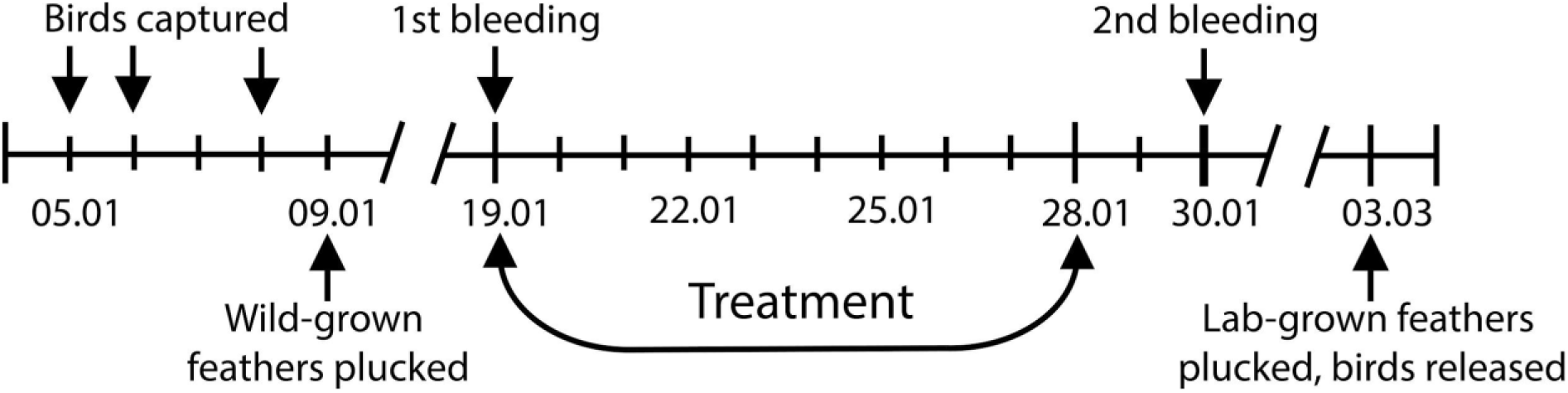
Timeline of the experiment. The experiment took place in 2015, dates are in DD/MM format. Wild-grown feathers were plucked on 09.01 and lab-grown feathers on 03.03. Birds were treated for 10 days (19.01-28.01) with anticoccidial drug Toltrazuril or with antibiotic Metronidazole. Birds were bled twice, before treatment on 19.01 and post treatment on 30.01. The duration of the study (09.01-03.03) was based on the time it takes greenfinches to regrow plucked feathers (Hõrak et al., 2013).

### Measurement of chroma

We plucked the left and right outermost wild-grown tail feathers (rectrices) on 9th January from 71 birds. Replacement feathers, grown during the study (lab-grown feathers) were collected on March 3rd, before the release of the birds. We placed collected feathers into a plastic bag and stored in the dark until measurements were carried out. Locations of color measurements on feathers are indicated in Fig. 2. Color was measured from the feathers placed on a black background, in an area of approximately 1 mm^2^, of the visible surface of the feather, using a spectrophotometer (Ocean Optics S2000) as described by Saks et al. (Saks et al., 2003). To estimate color, we calculated values of chroma (Endler, 1990). Chroma can be understood as a measure of the ‘purity’ or ‘saturation’ of a color, and has been shown to correlate with actual carotenoid concentration of feathers. Details for calculations of chroma are given in Saks et al. (Saks et al., 2003). The chroma measurements of left and right feather were repeatable in lab-grown feathers and in wild-grown feathers (r=0.7, F_60;61_=5.1, p<0.0001) (Lessells and Boag, 1987), however there was no correlation between chroma of wild-grown feathers and chroma of lab-crown feathers (r=0.093, p=0.46, N=65).

**Fig. 2.**
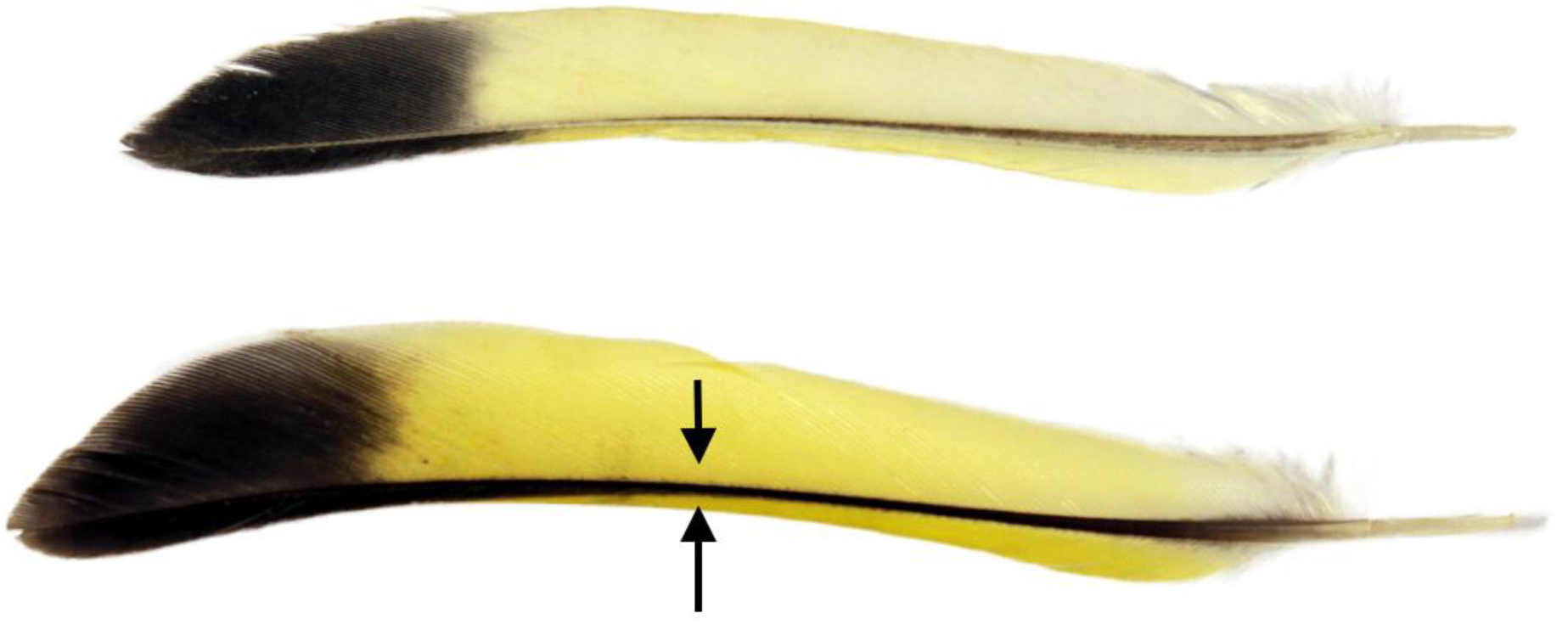
Measurement of feather chroma. The arrows indicate positions where chroma of yellow feather coloration was measured. Upper – lab-grown feather, lower – wild-grown feather.

### Infection intensity and blood analyses

Fecal samples for determination of coccidian (*Isospora* spp.) infection intensity were collected from all the birds on days 1, 14, and 20 as described by (Meitern, Lind et al. 2016) and all birds appeared naturally infected. Diagnosis of trichomonosis requires either sacrificing the birds or labour- and resource-intensive techniques (Amin et al., 2014). Therefore, we were not able to collect data on the infection status of trichomonads for this study, but previous investigations of dead birds have confirmed that trichomonosis is present in Estonian greenfinch population. For instance, three from the 46 wild-caught captive greenfinches that died in captivity in our lab in 2013 showed clear symptoms of trichomonosis, but the prevalence of sublethal levels of infection is not known (Männiste and Hõrak, 2014). The concentration of carotenoids was determined spectrophotometrically from 15μl of plasma, diluted in acetone as described elsewhere (Sild et al., 2011). The nutritional state was assessed based on plasma triglycerides, measured as described by Meitern, Lind et al. (2016). Plasma triglycerides increase during transport of fat to the adipose tissue and energy consuming organs and indicate amount of lipids absorbed and changes in fat stores (Jenni-Eiermann and Jenni, 1994).

### Statistics

The effect of experimental manipulation on feather chroma was analyzed with one-way ANOVA. The assumptions of parametric tests were met (normality of residuals, homogeneity of variances). We used Tukey’s post hoc test to determine differences between groups. The effect of experimental manipulation on plasma carotenoids was examined with repeated measures ANOVA, assuming that the effect of treatment will be revealed by a significant ‘time × treatment’ interaction term. All tests are two-tailed with a P-level below 0.05 as a criterion for significance. Sample sizes varied between analyses due to our inability to collect sufficient good quality blood samples from all birds. Analyses were performed with a package Statistica v. 12 (Statsoft Inc., Tulsa, OK). We examined only male greenfinches as carotenoid based sexual signal traits are more pronounced in males in this species and addition of another factor (sex) would have decreased test power. Female birds were used in a different study. We also explored correlation between plasma carotenoids, chroma and plasma triglycerides.

## Results

### Treatment’s effect on feather chroma

The experimental treatment had a significant effect on chroma of yellow parts of the lab-grown tail feathers (ANOVA F_2,62_=5.6; p=0.006) (Fig. 3). Birds that received METRO had higher chroma than in control group (Tukey’s post-hoc, p=0.012) and in TOLTRA group (Tukey’s posthoc p=0.016), whereas there was no difference in chroma between birds treated with TOLTRA and control group. There was no difference in chroma in the wild-grown feathers between the experimental groups (F_2,68_=0.43, p=0.70).

**Fig. 3.**
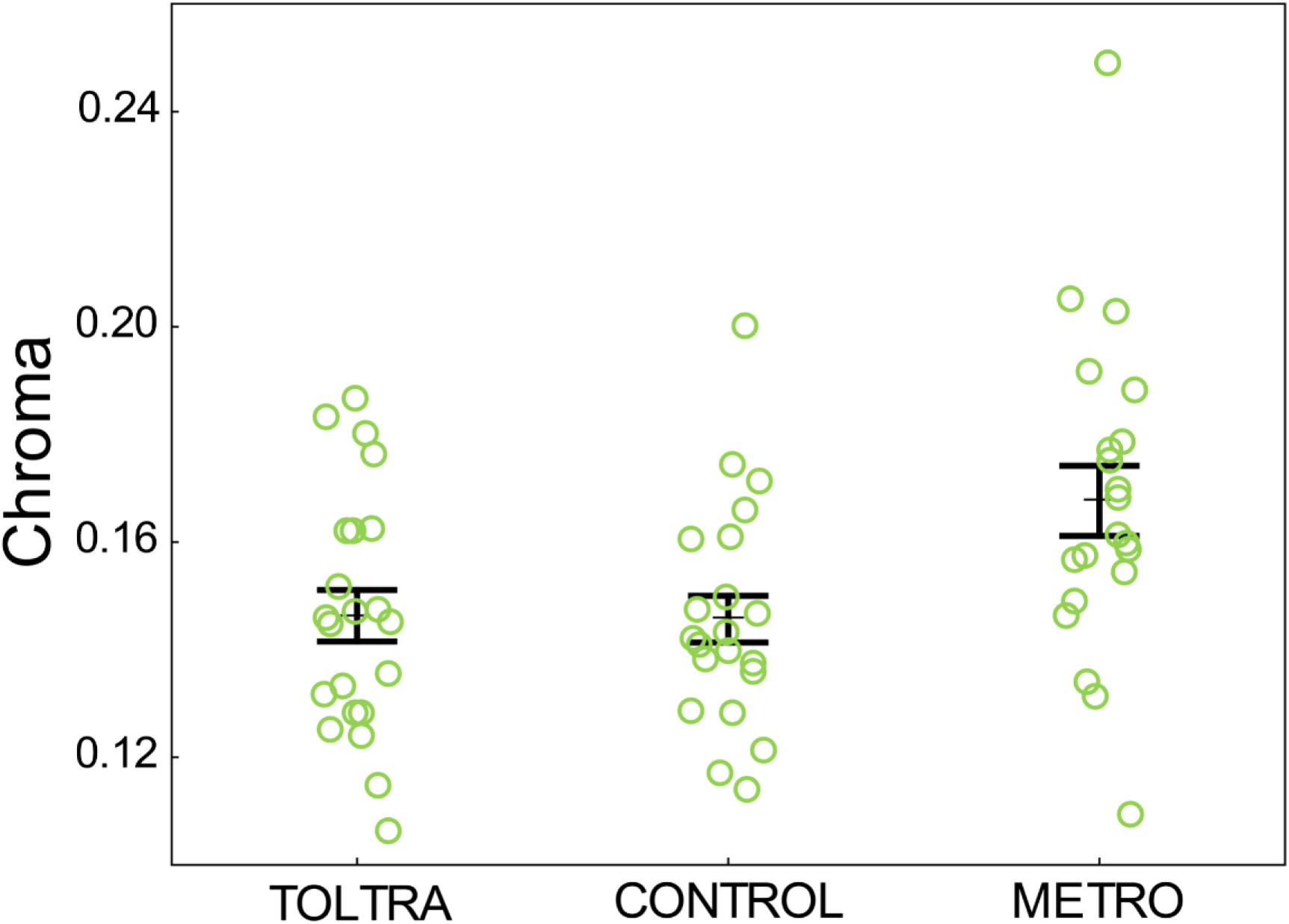
The effect of treatment on feather chroma. The treatment affected chroma of the yellow parts of the lab-grown feathers (N=63; ANOVA F_2,62_=5.6; p=0.006). Whiskers denote mean ± s.e.m. Birds that received metronidazole medication (N=20) had higher chroma than in control group N=21 (Tukey’s post-hoc, p=0.012) and in toltrazuril group (N=22) (Tukey’s post-hoc p=0.016).

### Infection intensity and plasma biomarkers

The effects of treatment on coccidian infection intensity are reported in previously published article on this study by R. Meitern and coauthors: medication with TOLTRA significantly reduced intensity of coccidian infection while METRO had no effect and did not differ from the control group (repeated measures ANOVA F=24.54,130 p<.00001) (figures in (Meitern et al., 2016).

The treatment had an effect on plasma carotenoids (repeated measures ANOVA F2,62=,5.02 p=0.0095). Plasma carotenoids declined in all groups, but the decline was steepest in the TOLTRA group. Carotenoids were significantly higher in TOLTRA group pre-treatment (ANOVA F2,67=3.73, p=0.03), but after medication there were no differences between groups (ANOVA F2,63=2.66, p=0.1). Plasma triglycerides were measured by (Meitern et al., 2016) and no significant effect of treatment on plasma triglycerides were detected (repeated measuresANOVA F2,47=1.71, p=0.19), but post-treatment, on 30^th^ January, plasma triglycerides were correlated positively with plasma carotenoids (r=0.41, N=58 p=0.0013). We also detected that chroma of lab-grown feathers and plasma carotenoids on 19^th^ January and 30^th^ January were correlated (respectively r=0.33, N=64, p=0.008 and r=0.42, N=62, p=0.0006). However, there was no correlation between plasma carotenoids and chroma of wild-grown feathers on neither of the days (r=0.12, N=70, p=0.34 and r=0.05, N=66, p=0.69).

### Discussion

Carotenoid coloration is suggested to signal parasite load and parasite resistance of the individual. Our goal was to test whether parasites have effect on yellow feather coloration through decreased nutrient acquisition. We administered anticoccidial drug TOLTRA or antibiotic METRO, that targets *Trichomonas* protozoa and anaerobic bacteria, to wild-caught captive greenfinches. We found that the treatment with METRO resulted in significantly higher chroma of yellow parts of the feathers, whereas feather color in birds who received anticoccidial TOLTRA did not differ from the control. There were no significant differences in plasma carotenoids nor in plasma triglycerides, indicating that treatment had no effect on absorption or availability of these nutrients. We suggest that treatment with antibiotic METRO attenuated negative effects of coinfection, which freed additional energetic resources that could be invested in ornamentation (del Cerro et al., 2010). Alternatively, METRO could improve vital cellular processes that also influenced modification and deposition of carotenoids to the feathers (Hill, 2011). Antibiotic METRO might have induced change in gut microbiota composition and some metabolites of microbiota, such as short-chain fatty acids (SCFAs), can affect mitochondrial function (Saint-Georges-Chaumet and Edeas, 2015, Franco-Obregón and Gilbert, 2017). There was no effect of treatment on yellow feather coloration in the TOLTRA group. One possible explanation could be that birds might have been naturally infected with mildly virulent strains of coccidia that had little effect on their health before the treatment. For instance, it has been demonstrated previously in captive greenfinches that coccidian strains infecting different individuals vary largely in their virulence and hosts differ in resistance (Hõrak et al., 2006).

One explanation why birds who received METRO grew yellower feathers might be that METRO had an effect on the coinfection dynamics that freed additional energy resources which could be allocated to production of ornaments. Most individuals in the wild are faced with coinfection of several parasites or several different strains of one parasite (Paterson, 2013) and indeed, all the greenfinches in our study were naturally infected with *Isospora* parasites. We suggest that the wide spectrum antibiotic METRO acted on *Trichomonas gallinae* and/or on multiple (bacterial) infections. Although we were not able to diagnose trichomonosis in birds used in this study, previous investigations of dead birds have confirmed that trichomonosis is present in Estonian greenfinch population (Männiste and Hõrak, 2014). The prevalence of Trichomonas parasites has been measured in the greenfinch population in Hesse, Germany where it showed high prevalence with 25% of the population being infected and the highly pathogenic strain was widespread (Quillfeldt et al., 2018). Coinfection can affect the virulence of the parasites (Kinnula et al., 2017), but also the host’s immune responses (Paterson, 2013). Israeli sparrows (*Passer domesticus biblicus*) that were simultaneously infected by Isospora and *Leucocytozoon gentili* experienced more severe symptoms of coccidiosis (Gill and Paperna, 2008). Carotenoid feather coloration was paler in blue tits (*Cyanistes caeruleus)* who were infected by different numbers of blood parasite genera compared to those infected just by one (del Cerro et al., 2010). Necrotic enteritis in poultry has been shown to occur when birds with preexisting coccidiosis are infected with pathogenic *Clostridium spp.* Strains (reviewed by (Shojadoost et al., 2012, Williams, 2005). Members of *Clostridium* genus are also sensitive to METRO as are many other anaerobic bacteria (Freeman et al., 1997). It is possible that by administering METRO, the birds were relieved from the adverse effects of coinfection. It has been suggested that simply expressing all the proteins that are necessary for carotenoid uptake, transportation, modification and deposition to ornaments can be energetically demanding itself (Hill, 2000). Therefore, reduction of available energy through allocating energy to fighting parasites or to other physiological functions can result in duller ornaments and indicate that carotenoids signal energetic state of the animal.

Antibiotics can affect host energy metabolism through altering gut microbiota composition. In addition to antiparasitic effects, antibiotics can be harmful as they can alter the normal microflora. However, recent research has found that some antibiotics can increase the abundance of beneficial bacteria (reviewed by (Ianiro et al., 2016). Administration of METRO to dogs resulted in an increase of beneficial bacteria in fecal microbiota, such as *Bifidobacterium, Enterococcus, Lactobacillus,* and *Streptococcus* (Igarashi et al., 2014) (but note that the study lacked control group and sample size was small). These bacteria can modulate the intestinal immune system and are important producers of short-chain fatty acids (SCFAs). SCFAs can be an energy source, but they also act as regulatory molecules that influence energy homeostasis and mitochondrial function (reviewed by (Saint-Georges-Chaumet and Edeas, 2015, Franco-Obregón and Gilbert, 2017). Many other bacterial metabolites (such as secondary bile acids, lipopolysaccharides, urolithins) have been shown to have regulatory and immunomodulatory effects (reviewed by (Heiss and Olofsson, 2018, Franco-Obregón and Gilbert, 2017). It is possible that although METRO treatment didn’t result in higher nutrient absorption, it might have elicited a shift in gut microbiota composition that improved host’s cellular processes.

We suggest that the effect of METRO on yellow feather coloration might have resulted from improved vital cellular processes that share pathways with carotenoid modification and carotenoid deposition into feathers. This is in accordance with the shared pathway hypothesis, which states that some core system functions pathways are shared with productions of ornamental traits (Hill, 2011). It has been proposed that modification of carotenoids depends on capacity to produce energy in the membranes of mitochondria (Hill, 2014). Mitochondria are target of many pathogens, but mitochondria also produce reactive oxygen species and are closely involved with immune pathways and signaling (reviewed by (Koch et al., 2017). According to endosymbiotic theory, the ancestors of mitochondria were once free-living prokaryotes that developed a symbiotic relationship with the eukaryotic ancestor cell, thus mitochondria shares several structural and functional features with the prokaryotes. For that reason, some metabolites of microbiota have profound effects on mitochondria (Saint-Georges-Chaumet and Edeas, 2015, Franco-Obregón and Gilbert, 2017) and therefore microbiota can affect mitochondrial function. Our results are consistent with the shared pathway hypothesis, however future experimental work on connections between mitochondrial function, ornamentation and microbiota are needed.

The lack of effect of TOLTRA on feather color might have been due to low virulence of natural infection and that the strains of coccidia were familiar to the birds. Greenfinches have better tolerance to their ‘own’ previously acquired parasites than to novel strains (Hõrak et al., 2006). Infection intensities were higher when birds were infected with strains that originated from several hosts, opposed to when birds were inoculated coccidia strains from one single host (Hõrak et al., 2006). That indicates variation of virulence in *Isospora* coccidia in greenfinches. We believe that the naturally occurring coccidian infection did not cause considerable intestinal damage in our study system that would have any effect on the absorption of carotenoids. It has been suggested that birds in the wild can tolerate *Isosporan* parasites well, but can experience more serious illness under stressful conditions (e.g captivity), poor sanitary conditions or with concurrent infections (Gill and Paperna, 2008, Swayne et al., 1991). Greenfinches however, have previously shown to tolerate captivity well based on low stress hormone levels in captivity (Lind et al., 2020) and several hematological indices (Sepp et al., 2010), thus the stress of captivity might not have been severe enough to induce exacerbation of the coccidian infection.

Lack of the treatment effect on plasma carotenoids and triglycerides suggests that there were no differences in carotenoid and triglyceride absorption between groups. Plasma carotenoid levels before the treatment were by chance significantly higher in birds who subsequently received TOLTRA. After the treatment plasma carotenoids did not differ between groups, however plasma carotenoid levels declined in all groups. We suggest that carotenoid levels in all the birds fell to the approximately same levels in the lab conditions because all birds received a constant amount of carotenoids. Negative effect of TOLTRA on plasma carotenoid levels cannot be ruled out, however, we are not aware of any such findings. It is thus more likely that equal carotenoid availability in the diet in captivity was responsible for uniform plasma carotenoid levels in all our treatment groups. Similar decrease in plasma carotenoid levels during captivity has been recorded in our previous experiments with greenfinches (Hõrak et al., 2004, Sild et al., 2011, Sepp et al., 2010).

To conclude, treatment of captive greenfinches with two antimicrobial agents – narrow spectrum anticoccidial Toltrazuril and a wide spectrum antibacterial/anti-trichomonad Metronidazole did not affect plasma carotenoid levels, indicating that neither of our treatments affected carotenoid absorption. Among the birds treated with Metronidazole, feathers grown during the experiment had significantly higher yellow chroma, which is indicative of higher deposition of dietary carotenoids into feathers. The latter finding provides indirect support to the hypotheses about the importance of microbiome for carotenoid metabolism, transportation, and/or deposition. Assuming that microbial metabolites can modulate mitochondrial and immune function by altering the efficiency of vital cellular processes, our findings are also consistent with the shared pathway hypothesis (Hill, 2011, Hill, 2014) whereby the mechanisms of production of ornaments share functional pathways with core-life supporting pathways. Specific mechanisms how microbiome relates to carotenoid metabolism and signaling thus await further investigation.

## Acknowledgements

We thank Elin Sild, Ulvi Karu, Richard Meitern, Janek Urvik and Liina Ots for help with bird maintenance and experiments.

